# A multiscale 3D chemotaxis assay reveals bacterial navigation mechanisms

**DOI:** 10.1101/2020.03.18.997841

**Authors:** Marianne Grognot, Katja M. Taute

**Affiliations:** Rowland Institute at Harvard University, 100 Edwin H. Land Blvd, Cambridge, MA 02142

## Abstract

How bacteria navigate environmental chemical gradients has implications ranging from health to climate science. The underlying behavioral mechanisms are unknown for most species due to a lack in techniques bridging scales from individual 3D motility behavior to the statistical power required to assess the resulting performance. We present the first demonstration of such a multiscale 3D chemotaxis assay and reveal that *Caulobacter crescentus* chemotaxis breaks with the *Escherichia coli* paradigm.

---

Chemotaxis enables bacteria to navigate external chemical fields and is now recognized as a key factor driving interactions of bacteria with their environment and each other, with wide-ranging effects that include promoting pathogenicity^1^, establishing symbioses^2^, and shaping geochemical fluxes^3^. Because individual bacterial motility behavior has a large random component, chemotaxis is usually assessed using population-level assays that average over the behavior of thousands of individuals, ranging from Adler’s classic capillary assay^4^ to modern approaches based on video microscopy^5^. While these approaches are highly sensitive and precise in detecting small chemotactic effects, they are blind to the underlying behavioral mechanisms. Berg’s pioneering 3D tracker capable of following a single bacterium swimming in 3D demonstrated the importance of resolving individual 3D motility behavior for revealing chemotactic mechanisms^6^, and the resulting understanding that *Escherichia coli* chemotaxis is controlled via the bias in the rotation direction of the flagellum^7^ has become a dominant paradigm in bacterial chemotaxis research^8^. Recent findings, however, suggest that many other species, including the majority of marine bacteria, may use a different strategy^9,10^, highlighting the need for efficient and broadly applicable methods of characterizing chemotactic behavior.

The pervasive inter-individual variability present even in genetically identical populations^11^ renders throughput and sampling crucial bottlenecks in characterizing chemotactic behavior based on trajectory data. Methods aimed at revealing potentially diverse chemotactic mechanisms must bridge the scales between individuals and populations by capturing motility behavior of individual bacteria with the statistical power to simultaneously reveal chemotactic performance, which is typically only accessible via ensemble averages. Here we introduce a novel chemotaxis assay that enables such a multiscale approach by harnessing a recently developed high-throughput 3D tracking method^12^ to capture individual bacterial behavior in the presence of microfluidically created chemical gradients for thousands of individuals within minutes. After validating our approach using the well-characterized *E. coli* model system, we demonstrate its power by revealing that the chemotactic mechanism of the fresh water bacterium *Caulobacter crescentus* breaks with the *E. coli* paradigm.

In our assay, typically 50-100 bacteria are tracked simultaneously in 3D in the center of a quasistatic, linear gradient field created in a microfluidic device (Fig. 1a, see Methods for details). Typically, thousands of trajectories are gathered in minutes, enabling a precise determination of the chemotactic drift velocity, v_d_, as the population-averaged velocity along the direction of the gradient (*x*) (Fig. 1b). We demonstrate the technique by assessing chemotaxis of the well characterized *E. coli* strain AW405 towards the non-metabolizable chemoattractant methyl aspartate (MeAsp, Fig.1b-f). Resolving the drift velocity, *v*_*d*_, as a function of vertical position, *z*, reveals that the drift velocity is roughly constant in the bulk liquid, but decreases sharply near the surfaces of the sample chamber (Fig. 1d). We attribute this decrease to trajectory curvature at the surfaces (see Fig. 1c) randomizing bacterial orientations and thus leading to a decreased chemotactic response. Such curvature results from hydrodynamic interactions with the surface^13^ and is thus present in common 2D motility assays that increase trajectory durations by either constraining the bacteria to a thin sample chamber^14^ or by limiting observations to a chamber surface^15^. The population-averaged drift velocity of 2.8±0.4 µm/s (mean ± sd) for bulk trajectories in a linear gradient of 10 µM/mm MeAsp aligns well with previously reported values of the chemotactic sensitivity (Supplementary Discussion), confirming that our technique provides an accurate quantification of chemotactic performance. In addition, our findings indicate that many standard 2D chemotaxis assays may be compounded by surface effects in their ability to quantify chemotactic performance, demonstrating the value of 3D tracking.

**Fig. 1:**
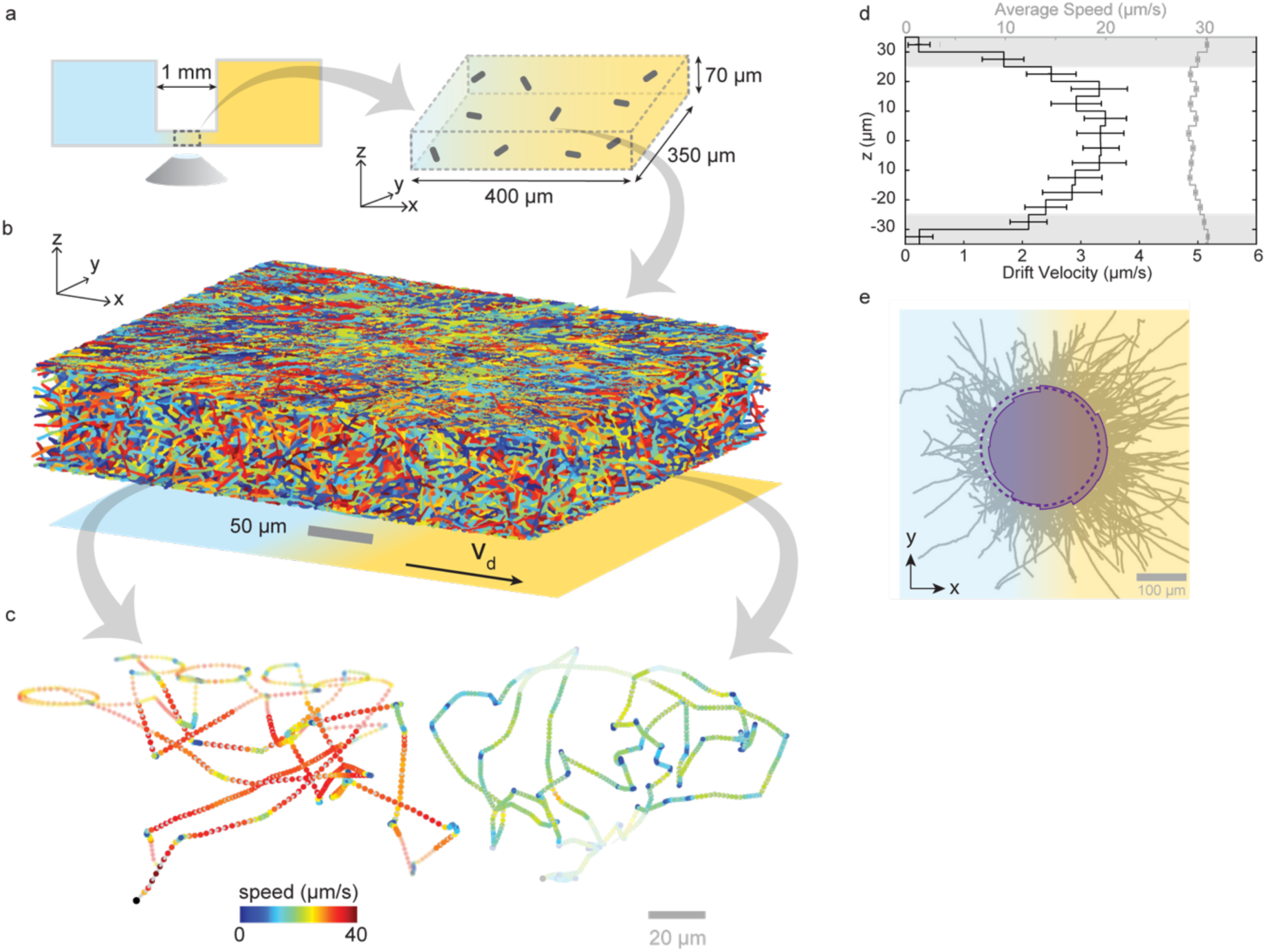
Schematic of multiscale chemotaxis assay and its application to E. coli strain AW405. a) A quasi-static linear chemical gradient is established between two reservoirs containing a uniform concentration of bacteria. Bacteria are observed in the central portion of the linear gradient. b) 5,045 individual trajectories with a minimum duration of 5 frames and containing 37,080 s of total trajectory time, obtained in 9 min of recording at 15 Hz in a typical experiment. c) Two example trajectories (durations 63 s and 65 s) showing run-tumble motility in bulk solution and circular segments near the chamber surface (within 10 µm distance, shaded). d) Drift velocity (black, defined as the average speed along the gradient direction, x) and average swimming speed (grey) as a function of height, z. Only bulk trajectories (covering 35 % of total trajectory time, and defined as trajectory segments with a distance of more than 10 µm to the surface) are retained for further analysis. Error bars reflect standard errors of the mean. e) Bulk trajectories aligned with the origin (grey) and polar pdf of instantaneous swimming directions projected in the x-y plane (aqua, solid line). A flat distribution (dashed) is shown for reference.

To demonstrate the power of our approach for revealing novel chemotactic behaviors, we turn to the fresh water bacterium *C. crescentus* whose weak, cell cycle-dependent chemotaxis response^16,17^ has been challenging to capture. Its “run-reverse-flick” motility^9,18^ is driven by a single polar flagellum. Either direction of flagellar rotation results in locomotion, with the flagellum either pushing the cell body forward or pulling it backward. Reversals (turns by ∼180°) occur with the transition from pushing to pulling, whereas the reverse transition is accompanied by a turn about a smaller angle (“flick”, Fig. 2a).

**Fig. 2:**
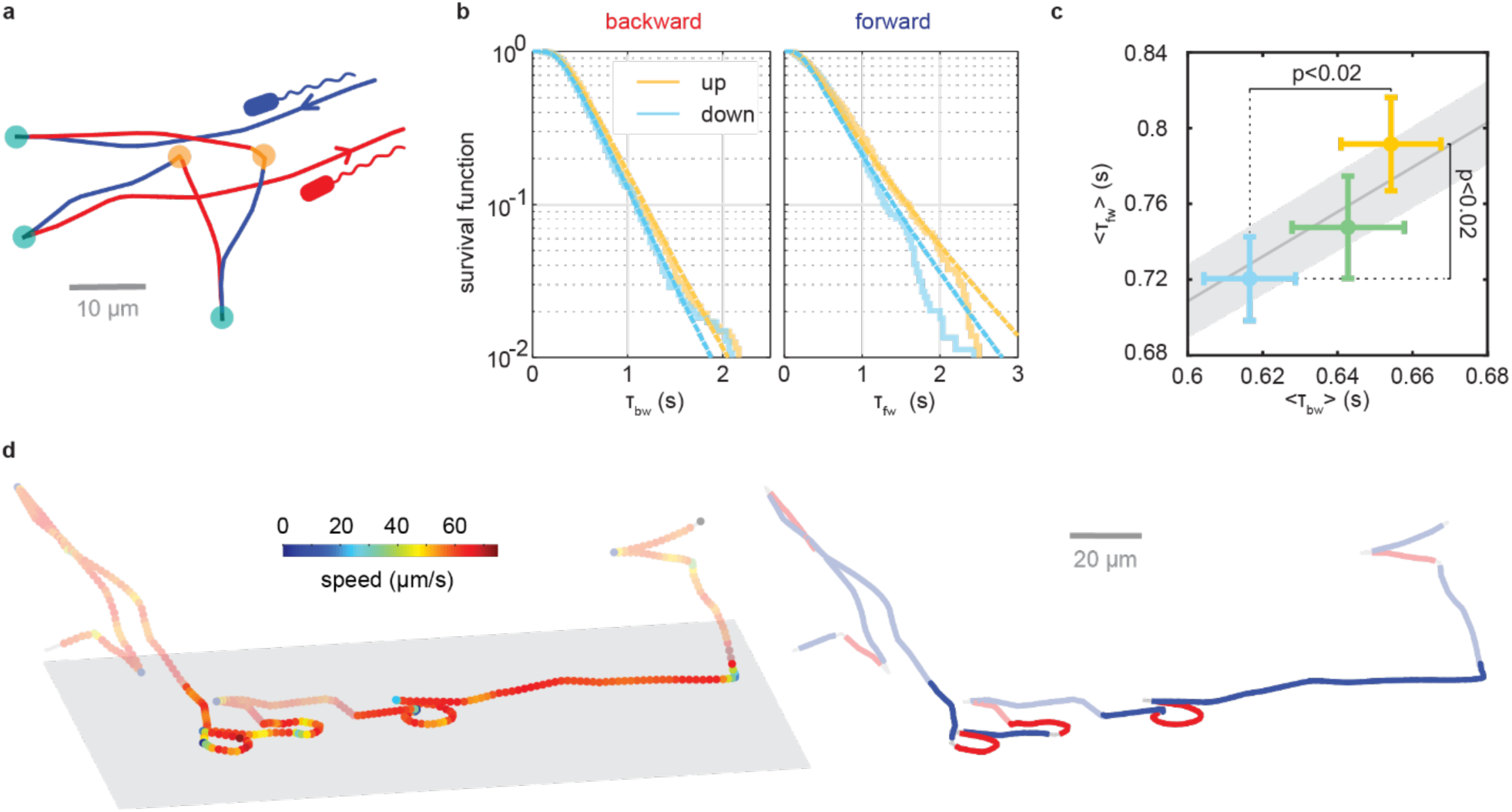
C. crescentus chemotaxis. a) Example trajectory showing alternating backward (red) and forward (blue) runs, separated by reversals (aqua) or flicks (orange). b) Survival function (fraction of events above threshold) for forward (left panel) and backward (right panel) runs, going up (yellow) or down (blue) the gradient, defined as falling within a 36° cone around the positive or negative x-axis, respectively. The dashed lines show corresponding maximum likelihood inverse Gaussian distribution functions. c) Average run durations up (yellow), down (blue) or perpendicular to (green) the gradient (defined by 36° cones around positive x-axis, negative x-axis, or y-axis, respectively), and p-values for one-sided t-tests between durations up and down the gradient. Error bars reflect standard error of the mean. d) Example trajectory color-coded by speed (left) and run direction (right, red: backward, blue: forward), with near-surface segments shown at higher saturation than bulk segments.

We recorded 79,244 individual 3D bulk trajectories of motile *C. crescentus* cells navigating a 1 mM/mm xylose gradient (see Supplementary Methods and Supplementary Table 2 for details). Strikingly, 54% of the more than 123,000 s of total trajectory time we obtained consists of runs with no turns and no discernible chemotactic drift up the gradient. In *C. crescentus*’s life cycle, cell division occurs in surface-bound sessile cells and produces motile swarmer cells whose transition back to sessile cells is accompanied by the degradation of chemoreceptors^17^ which are likely required for turning. We thus speculate that the smooth-swimming subpopulation may be in the early stages of the transition. Such straight swimming is likely to accelerate the rate of surface encounters and may be an integral part of *C. crescentus*’s strategy for completing the swarmer-to-sessile cell transition.

The chemotactic drift speed of the remaining, turning population amounts to only 0.26±0.12 µm/s (mean ± SEM), corresponding to less than 0.5% of their average swimming speed of 56 µm/s. To our knowledge, this is the first measurement of a chemotactic drift speed in *C. crescentus*, demonstrating the sensitivity enabled by the high throughput of our technique. The speed of backward runs, with the flagellum pulling the cell, is 2.1±0.1% higher than that of forward ones (Supplementary Fig 2b.). Forward and backward run duration distributions are approximated well by inverse Gaussian distributions (Fig. 2b), consistent with previous reports^15,19^.

*C. crescentus* chemotaxis has been assumed to follow the *E. coli* paradigm where the cytoplasmic concentration of phosphorylated CheY ([CheY-P]), the chemotaxis response regulator, determines the bias in the direction of motor rotation^19^. In *E. coli*, one direction of flagellar rotation supports locomotion (“runs”), whereas the other direction induces reorientation events (“tumbles”). Chemotaxis is achieved by dynamic modification of the bias in rotation direction so as to increase the duration of runs aligned with the gradient direction. Early studies hypothesized that in *C. crescentus*, forward runs driven by clockwise (CW) rotation correspond to *E. coli* runs, while backward runs driven by counterclockwise (CCW) rotation are equivalent to *E. coli* tumbles because of their shorter duration^20,21^. Given similar forward and backward swimming speeds, a bias towards CW rotation would then yield net displacements in the forward swimming direction. In line with this hypothesis, a recent study of 2D surface swimming behavior in oxygen gradients found that forward, but not backward runs, are extended when directed up the gradient compared to down^15^.

In contrast with this hypothesis, we find that both forward and backward run segments are extended when ascending, versus descending, a chemoattractant gradient (Fig. 2b,c). In fact, our data support a constant motor bias, that is, a constant ratio of forward vs backward swimming interval durations (Fig. 2c) and thus indicate a radical break with the *E. coli* paradigm in *C. crescentus* chemotaxis. This interpretation is also consistent with the puzzling previous observation that, in sharp contrast to *E. coli*^*11*^, *C. crescentus* shows hardly any variability in motor bias between individuals^19^, suggesting that its motor bias might be unaffected by cytoplasmic fluctuations in [CheY-P].

The apparent conflict with previous findings^15^ likely results from a technical artefact imposed by the constraints of 2D tracking: to increase the typical time a bacterium spends in the focal plane, 2D bacterial tracking had been performed at the sample chamber surface which hydrodynamically attracts the bacteria^22^. The surface-induced trajectory curvature is more pronounced in the backward than in the forward runs^15,23^ (Fig. 2d), thus likely diminishing the chemotactic response more strongly for backward than for forward runs. This example highlights the crucial significance of full 3D behavioral information when assessing chemotactic mechanisms: 3D tracking enables the acquisition of long trajectories without a need for bacterial confinement as well as accurate turning angle measurements for faithfully determining bacterial orientation even for short trajectories with few turning events.

The mechanism we unveil for the alpha proteobacterium *C. crescentus* aligns with recent findings that also the singly flagellated gamma proteobacteria *Vibrio alginolyticus*^9^ and *Pseudomonas aeruginosa*^10^ extend both forward and backward swimming intervals during chemotaxis and suggests that this mechanism may be much more common than previously assumed. To our knowledge, no polarly flagellated species has conclusively been shown to follow the *E. coli* scheme, raising the intriguing possibility that the influential *E. coli* paradigm may reflect a special case limited to peritrichously flagellated bacteria.

In summary, our multiscale technique offers unprecedented, simultaneous access to individual and population level 3D motility behavior and is poised to offer unique insights into novel chemotactic mechanisms as well as the effects of phenotypic heterogeneity on population-level motility behaviors. In contrast to many flow-based chemotaxis assays^24^ that are limited to liquid environments, our assay is compatible with environments such as hydrogels that more closely mimic the complexities of many natural habitats, and thus paves the way for studies of chemotactic mechanisms in ecologically relevant settings.

## Supplementary Information

### Supplementary Discussion

#### Consistency of measured *E. coli* drift speed with literature values

Very few methods allow direct measurements of the drift velocity. Colin et al^5^ used a highly sensitive population scale approach to determine a drift velocity v_d_ normalized by the product of motile fraction α and average speed v_m_ for AW405 of v_d_/αv_m_ ∼0.08-0.09 for a 10 µM/mm gradient, which translates in our case (α=1, v_m_=29.9 µm/s) to an expected drift of 2.4-2.7 µm/s, in excellent agreement with our result of 2.8(±0.4) µm/s. Recently, Schauer et al.^25^ presented drift velocities directly computed from 2D trajectory data for the much slower swimming strain MG1655 (mean speed 16 µm/s). In addition to the strain difference, these results cannot be compared quantitatively to ours because the 2D approach introduces biases that are hard to correct^25^. In particular, mandating a minimum trajectory duration in 2D limits the analyzed set of bacteria to those that move within the focal plane, which results in an over-estimate of the drift velocity as it discriminates against bacteria that move perpendicular to the gradient.

Other publications refer to the chemotactic sensitivity. Ahmed et al.^26^ infer the drift velocity from the population average swimming speed and the measured asymmetry in the time spent swimming up versus down the gradient for various gradient conditions, and then use the following relation to determine the chemotactic sensitivity:

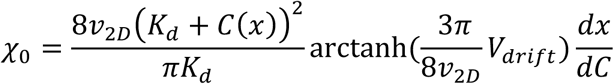

where 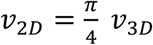 is the average bacterial swimming speed, *K*_*d*_ is the receptor/ligand dissociation constant, *V*_drift_ is the chemotactic drift, *C* is the MeAsp concentration and *x* the position in the gradient. Using this relation with 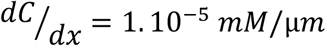, *C*(*x*) = 5 μ*M*, and *K*_*d*_ = 0.125 *mM*^27^, we obtain *χ*_0_ = 11(±2) 10^−4^ cm^2^/s from our drift velocity value of 2.8(±0.4) µm/s, which aligns very well with previously reported values for the same strain (2.4.10^−4^ cm^2^/s^28^, 5 .10^-4^ cm^2^/s^29^, 12.4 .10^-4^ cm^2^/s^26^).

**Supplementary Figure 1:**
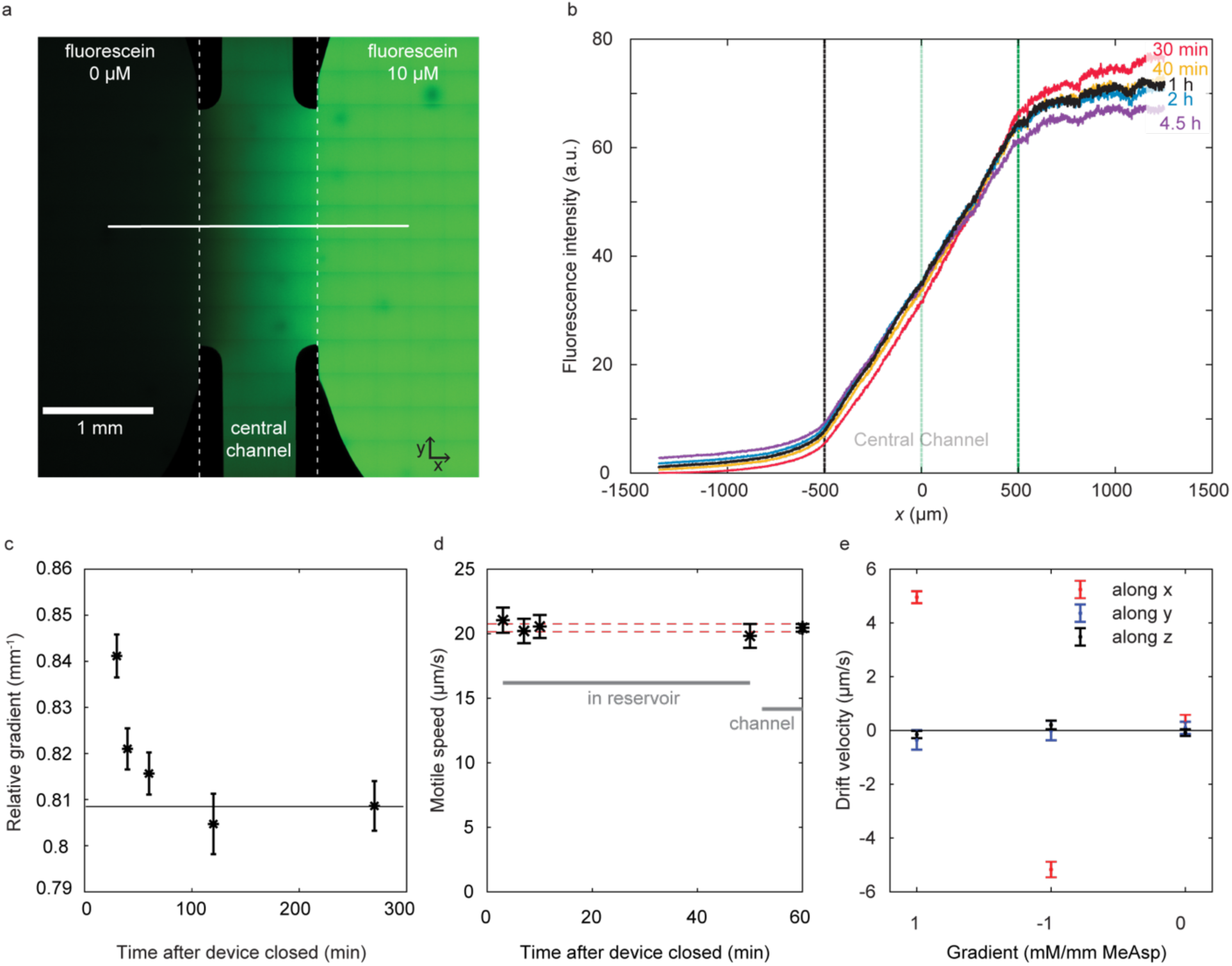
a) Top view of chemotaxis chamber with fluorescein gradient, stitched from tiled confocal fluorescence images. The gradient is established in a 1mm long central channel between two large reservoirs (left and right). b) Temporal establishment and stability of the fluorescein gradient assessed by confocal fluorescence imaging along a line in the center of the channel. c) Relative gradient in the center of the device as a function of time after closing the device, normalized relative to the intensity in the right reservoir. d) Individual swimming speed distribution of the motile population (defined as having a mean swimming speed larger than 10 µm/s) over time after closing the device. e) Average bacterial velocities measured in all 3 dimensions in the presence and absence of a 1mM/mm methyl aspartate gradient, and when the device is rotated by 180° about the z axis. The magnitude of the measured drift up the gradient changes by less than 1% when the chamber is flipped. All error bars are standard errors of the mean.

**Supplementary Figure 2:**
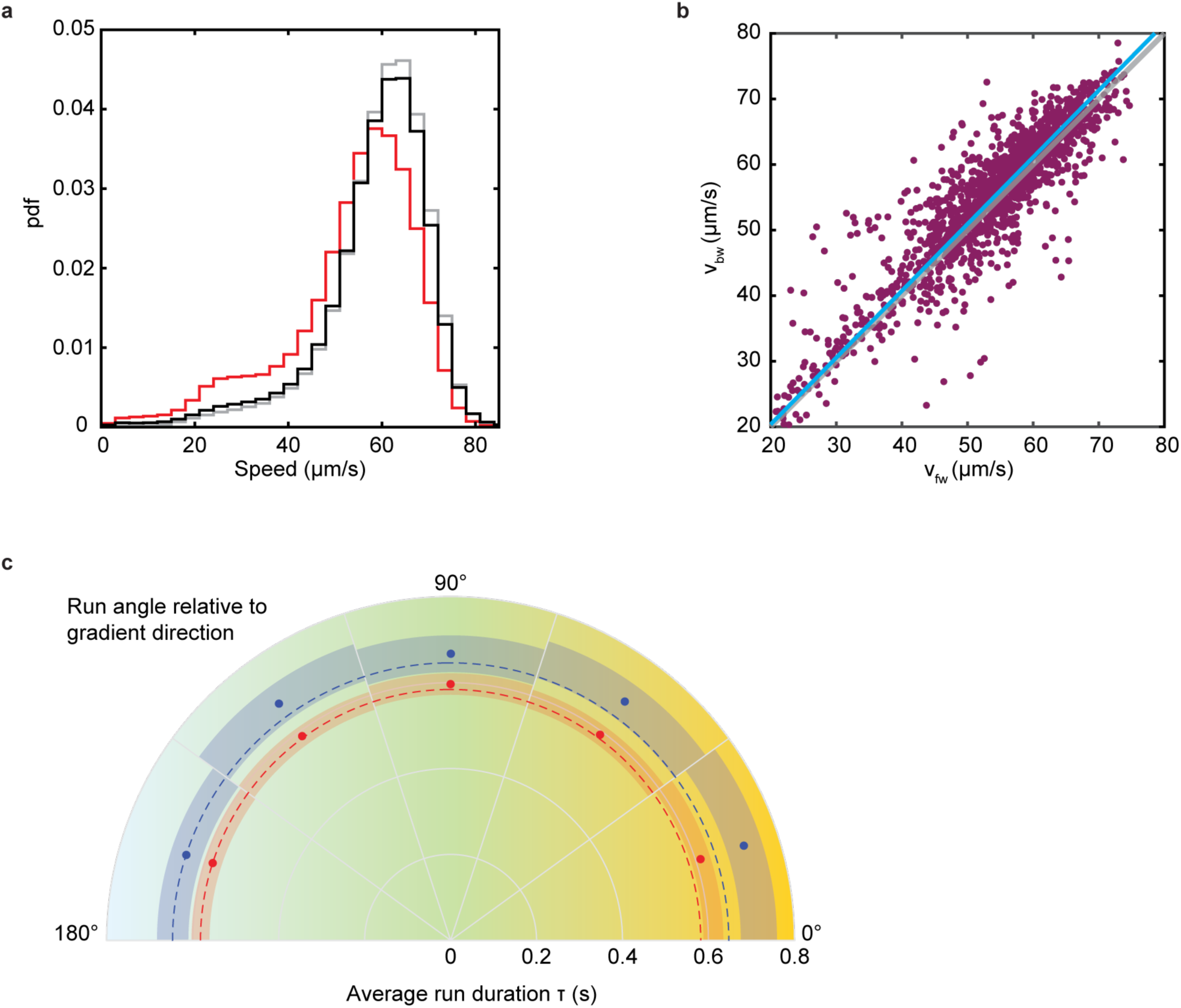
a) Instantaneous swimming speed distributions for the full motile bulk population (black, set 2 in Supplementary Table 2), the smooth-swimming population (grey, set 4) and that retained for run analysis (red, set 7), containing all trajectories with at least one run of defined duration and bacterial orientation. We attribute the subtle deviation in average swimming speed to selection bias, rather than biological differences. Conditions that are more likely to be met by longer trajectories select for bacteria of lower speed because they spend more time in the field of view. b) Backward and forward swimming speed (*v*_*bw*_ and *v*_*fw*_, respectively) in the same individuals reveal that backward runs are approximately 2.1% faster. Cyan: orthogonal linear fit with zero intercept, weighted by the number of contributing runs per individual (grey: unity, for comparison). c) Radial plot of average run durations as a function of projected angle to the x-axis in the x-y plane. Shading indicates 95% confidence intervals.

## Supplementary Methods

### Microfluidic device and gradient stability

Quasi-static chemical gradients are created in a commercially available microfluidic device (IBIDI µ-slide Chemotaxis) featuring an approximately 70 µm high, 1mm long, and 2 mm wide channel connecting two 65 µl reservoirs. Gradient establishment and stability over time were characterized for fluorescein gradients (10µM/mm in MotM, see Supplementary Table 1 for media compositions) using a confocal microscope (Zeiss AXIO Imager.Z2). The 488nm line was focused through a 20x water immersion objective and fluorescence emission collected in the 500–585 nm window with a pinhole adjusted to 30 µm diameter. A 4.5 h time series of line scans in the gradient direction across the center of the device was acquired after closing the device. A large scale 4×4 mm view of the device was obtained by tiling 2D scans (Supplementary Fig. 1a) acquired 5h after closing the device. The gradient is established within minutes of filling the device and shows a deviation of less than 4% from the final plateau value after 30 minutes (Supplementary. Fig. 1c). No detectable variation in relative gradient magnitude is observed in the time range from 40 min to 4.5h after filling the device (Supplementary Fig. 1c).

### E. coli experiments

#### Bacterial culturing

Overnight cultures were inoculated from a frozen glycerol stock of *E. coli* AW405 (a kind gift of Howard Berg) in 2 ml TB and grown to saturation at 30°C, 250 rpm. Day cultures were inoculated with the overnight cultures at 1:200 dilution in 10 ml TB and grown at 33.5°C, 250 rpm, until they reached an OD between 0.3 and 0.35 at 600nm. Volumes of 1 ml of bacterial culture were washed by three rounds of centrifugation in 1.5 ml Eppendorf tubes (6 min at 2,000 rcf), each followed by gentle resuspension in 1 ml of motility medium MotM. They were diluted to a target OD of 0.003 (for acquisitions in a gradient) or 0.005 (no gradient) in MotM supplemented with 0.002% Tween-20, with or without chemoattractant, for injection into the chemotaxis device.

#### Sample preparation

The device’s reservoirs were filled with the two bacterial solutions (with or without chemoattractant) following a modified version of the manufacturer’s “Fast Method” protocol. First, the entire device was overfilled with buffer free of chemoattractant or bacteria through the filling ports, and then the central channel’s ports were closed with plugs. 65 µl was removed from one reservoir, replaced by 65 µl of chemoattractant-free bacterial solution, and then this reservoir’s ports were closed. Finally, all liquid was removed from the other reservoir and replaced with bacterial solution containing chemoattractant. Key to reproducible gradients is to not overfill this reservoir to avoid liquid flow in the central channel when the last two ports are closed. A uniform bacterial density across the device ensures that any population drift observed is not the result of a diffusive flux, but likely indicates chemotaxis. For control measurements, neither bacterial solution contained chemoattractant. At the bacterial densities used (OD of 0.005 or less), oxygen depletion is unlikely, and we do not observe a change in *E. coli* swimming speed in the reservoirs over the course of 1 h (Supplementary Fig. 1d).

#### Data acquisition

Phase contrast microscopy recordings were obtained on a Nikon Ti-E inverted microscope using an sCMOS camera (PCO Edge 4.2) and a 40x objective lens (Nikon CFI SPlan Fluor ELWD 40x ADM Ph2, correction collar set to 1.2 mm to induce spherical aberrations^12^) focused at the center of the channel in all three dimensions. 3D bacterial trajectories were extracted^12^ for a field of view of ∼350 µm x 300 µm laterally (x, y) and over the entire depth (z) of the channel for typically several dozen individuals at a time. For *E. coli*, recordings were obtained starting from 50 minutes after filling the device. Three 3-min recordings were obtained at 15 fps.

#### In-device conditions

We confirmed that the *E. coli* motile population we tracked in the central channel was representative of the whole population by also acquiring trajectories in one chamber during an experiment. The average speed and population distributions were not significantly different, except for the reduced non-motile population fraction in the central channel. The average speed of the population was also monitored and no difference between 3 minutes and 1 hour after closing device was noted (Supplementary Fig. 1d).

#### Data analysis

3D Trajectories were extracted from phase contrast recordings using a high-throughput 3D tracking method based on image similarity between bacteria and a reference library^12^. Positions were smoothed using 2^nd^ order ADMM-based trend-filtering^30^ with regularization parameter *λ* = 1, and speeds computed as forward differences in positions divided by the time interval between frames. All trajectories with an average speed below a threshold were considered non-motile and discarded. The threshold was set at 15 µm/s unless noted otherwise. The *z* position of the top and bottom of the chamber were identified by visual inspection of trajectory data. All trajectory segments within 10 µm of the top or bottom of the central channel were removed to avoid surface interaction effects. We combine data from three biological repeats each yielding three recordings. The drift velocity is the average of the *x* component of all 3D speed vectors from all bacteria. Across 3 biological repeats, we obtain an absolute drift velocity of 2.8(±0.4) µm/s (mean ± sd) for *E. coli* in a 10 µM/mm MeAsp gradient, and observe no chemotactic drift along either the *y* or *z* axis (0.1±0.3 µm/s and 0.09±0.1 µm/s, respectively). A control chamber without a gradient showed no drift either (0.34±0.8 µm/s along *x*). For data obtained from a single experiment (Supplementary Fig. 1e), we estimate the noise on the drift measurement by jackknifing data into sets of 150 trajectories, and computing a standard error. For drift as a function of *z*, trajectories are first sliced into segments by *z* bin, and jackknifing is performed for each *z* bin.

### C. crescentus experiments

#### Bacterial culturing and sample preparation

Overnight cultures were inoculated from individual *C. crescentus* (CB15, ATCC 19089) colonies, grown on 1.5% agar PYE plates streaked from glycerol stock, and grown to saturation in 2ml PYE at 30°C, 200 rpm. Day cultures were inoculated at a dilution of 1:20 (v/v) in M2G^31^ or PYE and grown to an OD600 of at least 0.3 (a few hours for PYE, around 12 hours for M2G). Because the cell-cycle dependent motility of *C. crescentus* is quickly lost, we opted to grow the cells directly inside the device and let them produce swarmer cells while the gradient is being established. To this end, the day culture was again diluted 1:1 in fresh medium and injected into both reservoirs of the device. When the chamber walls were colonized by a sufficient density of stalked cells as determined by visual inspection under the microscope (a few hours for PYE, 2 days for M2G), the device was rinsed several times with fresh M2G, until no swimming bacteria were observed. Then fresh M2G and 1mM xylose / M2G, respectively, were injected into the reservoirs to create a 1mM/mm xylose gradient in the central channel.

#### Data acquisition

Because of the cell cycle-dependent chemotaxis^16^ of *C. crescentus*, we favored acquiring data as early as possible over waiting for perfect gradient stability. Recordings spanning 2.5 minutes at 30 fps were acquired from 20 to 55 minutes after closing the device, in five biologically independent experiments, totaling a cumulated acquisition time of 75 minutes.

#### Data analysis

Trajectories were obtained and smoothed as for *E. coli*, except for the ADMM regularization parameter being set to *λ* = 0.3. To account for the layer of attached cells lining the surface, only segments with a distance of more than 13 µm from the top or bottom chamber surface were retained to avoid surface interaction effects. Supplementary Table 2 details statistical characteristics of the subsets of data used for analysis. Standard errors on drift velocities are obtained by jackknifing as described for *E. coli*.

#### Run-reverse-flick analysis

The turning event detection is based on the local rate of angular change, computed from the dot product between the sums of the two consecutive velocity vectors preceding and subsequent to a time point. The threshold for a turn to begin is an α-fold rate relative to the median rate of angular change rate of the run segments, as determined in three iterations of the procedure. We determined by visual inspection of trajectories that a factor α=6 gave satisfactory results. A new run begins with at least two time points (at least 0.066 s) under this threshold. Backward (CCW rotation) and forward (CW rotation) runs were identified as runs with a turning event under 130°, respectively at the end or at the beginning the run, and a turning event above 150° at the other end of the run. A total of 5,342 backward and 2,898 forward runs were identified within a subpopulation of 6,230 trajectories.

---

**Supplementary Table 1:**
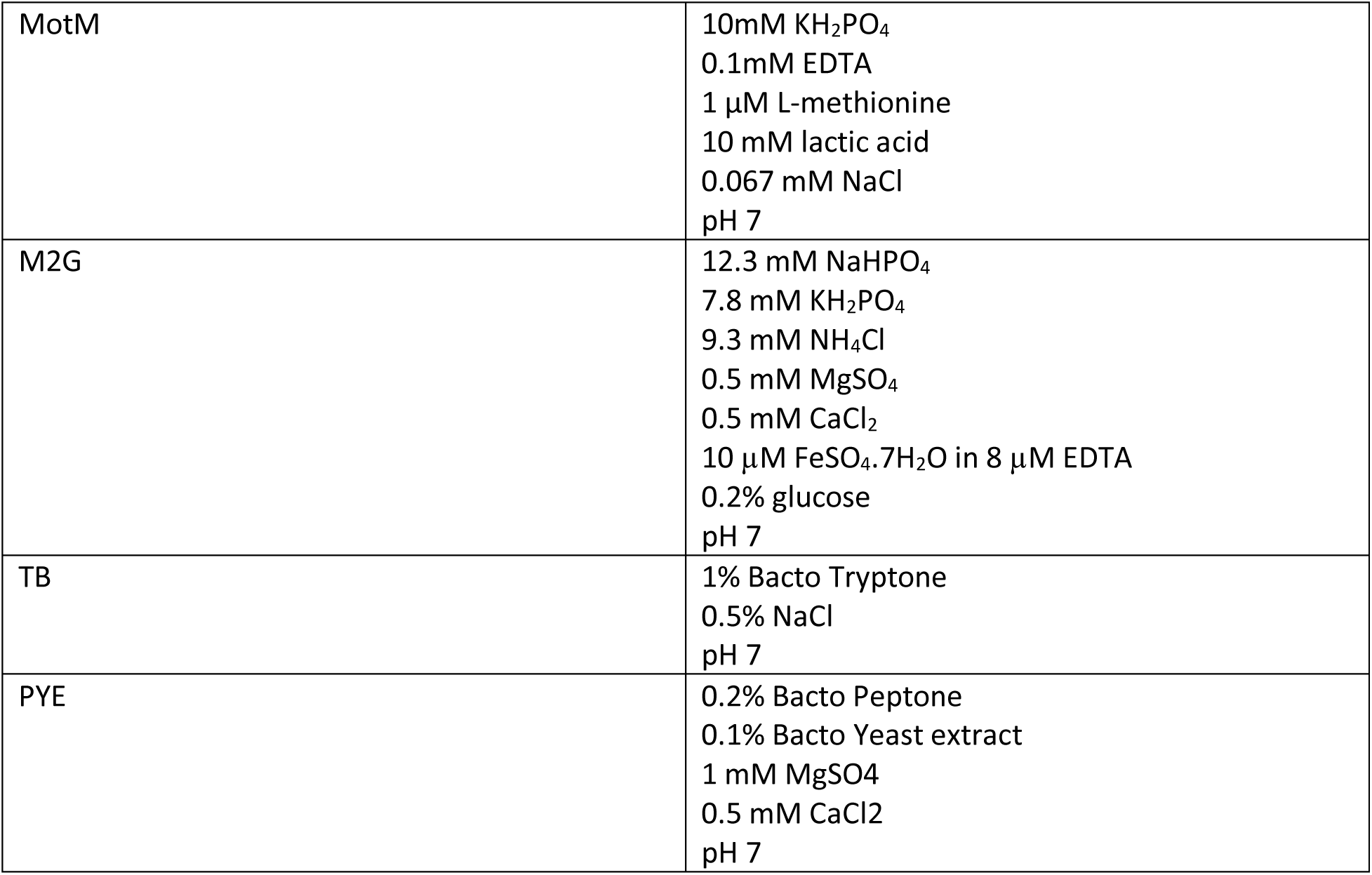
Growth and motility media used.

|  |  |
| --- | --- |
| MotM | 10mM $\text{KH}_2\text{PO}_4$<br>0.1mM EDTA<br>1 $\mu\text{M}$ L-methionine<br>10 mM lactic acid<br>0.067 mM NaCl<br>pH 7 |
| M2G | 12.3 mM $\text{NaHPO}_4$<br>7.8 mM $\text{KH}_2\text{PO}_4$<br>9.3 mM $\text{NH}_4\text{Cl}$<br>0.5 mM $\text{MgSO}_4$<br>0.5 mM $\text{CaCl}_2$<br>10 $\mu\text{M}$ $\text{FeSO}_4 \cdot 7\text{H}_2\text{O}$ in 8 $\mu\text{M}$ EDTA<br>0.2% glucose<br>pH 7 |
| TB | 1% Bacto Tryptone<br>0.5% NaCl<br>pH 7 |
| PYE | 0.2% Bacto Peptone<br>0.1% Bacto Yeast extract<br>1 mM $\text{MgSO}_4$<br>0.5 mM $\text{CaCl}_2$<br>pH 7 |

**Supplementary Table 2:**
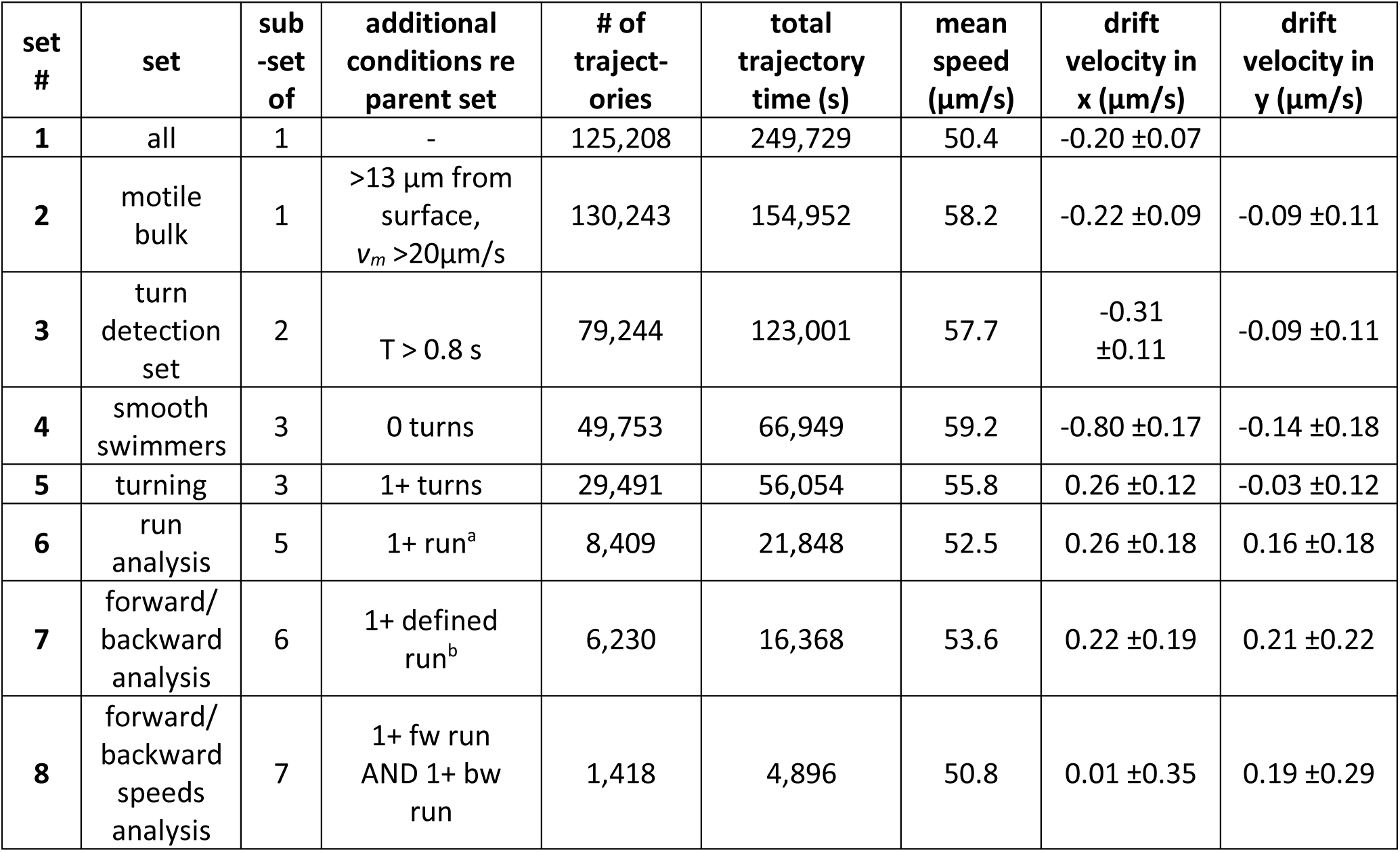
Definitions and properties of subsets of *C. crescentus* trajectories used for analysis. *v*_*m*_: mean swimming speed, *T*: trajectory duration. ^a^ A run is included here if its beginning and end are detected. ^b^ Defined runs are those whose bacterial orientation can be determined based on the magnitude of the preceding and subsequent turns.

## References

1 Matilla, M. A. & Krell, T. FEMS Microbiology Reviews 42, 202, (2018).

2 Raina, J.-B., Fernandez, V., Lambert, B., Stocker, R. & Seymour, J. R. Nature Reviews Microbiology 292, 1096, (2019).

3 Seymour, J. R., Amin, S. A., Raina, J.-B. & Stocker, R. Nature Microbiology 2, 17065, (2017).

4 Adler, J. Journal of General Microbiology 74, 77–91, (1973).

5 Colin, R., Zhang, R. & Wilson, L. G. Journal of The Royal Society Interface 11, 20140486–20140486, (2014).

6 Berg, H. C. & Brown, D. A. Nature 239, 500–504, (1972).

7 Larsen, S. H., Reader, R. W., Kort, E. N., Tso, W.-w. & Adler, J. Nature 249, 74–77, (1974).

8 Webre, D. J., Wolanin, P. M. & Stock, J. B. Current Biology 13, R47–49, (2003).

9 Xie, L., Altindal, T., Chattopadhyay, S. & Wu, X.-l. Proceedings of the National Academy of Sciences 108, 2246–2251, (2011).

10 Cai, Q., Li, Z., Ouyang, Q., Luo, C. & Gordon, V. D. mBio 7, e00013, (2016).

11 Spudich, J. L. & Koshland, D. E. Nature 262, 467–471, (1976).

12 Taute, K. M., Gude, S., Tans, S. J. & Shimizu, T. S. Nature Communications 6, 1–9, (2015).

13 Lauga, E., DiLuzio, W. R., Whitesides, G. M. & Stone, H. a. Biophysical Journal 90, 400–412, (2006).

14 Waite, A. J., Frankel, N. W., Dufour, Y. S., Johnston, J. F., Long, J. & Emonet, T. Molecular Systems Biology 12, 895–814, (2016).

15 Morse, M., Colin, R., Wilson, L. G. & Tang, J. X. Biophysical Journal 110, 2076–2084, (2016).

16 Laub, M. T., McAdams, H. H., Feldblyum, T., Fraser, C. M. & Shapiro, L. Science 290, 2144–2148, (2000).

17 Alley, M. R. Maddock & Shapiro, L. Science 259, 1754–1757, (1993).

18 Liu, B., Gulino, M., Morse, M., Tang, J. X., Powers, T. R. & Breuer, K. S. Proceedings of the National Academy of Sciences 111, 11252–11256, (2014).

19 Morse, M., Bell, J., Li, G. & Tang, J. X. Physical Review Letters 115, 312, (2015).

20 Ely, B., Gerardot, C. J., Fleming, D. L., Gomes, S. L., Frederikse, P. & Shapiro, L. Genetics 114, 717–730, (1986).

21 Koyasu, S. & Shirakihara, Y. Journal of Molecular Biology 173, 125–130, (1984).

22 Berke, A. P., Turner, L., Berg, H. C. & Lauga, E. Physical Review Letters 101, 038102–038104, (2008).

23 Magariyama, Y., Ichiba, M., Nakata, K., Baba, K., Ohtani, T., Kudo, S. & Goto, T. Biophysical Journal 88, 3648–3658, (2005).

24 Mao, H., Cremer, P. S. & Manson, M. D. Proceedings of the National Academy of Sciences 100, 5449–5454, (2003).

## References

12 Taute, K. M., Gude, S., Tans, S. J. & Shimizu, T. S. Nature Communications 6, 1–9, (2015).

25 Schauer, O., Mostaghaci, B., Colin, R., Hürtgen, D., Kraus, D., Sitti, M. & Sourjik, V. Scientific Reports 8, 9801, (2018).

26 Ahmed, T. & Stocker, R. Biophysical Journal 95, 4481–4493, (2008).

27 Mesibov, R., Ordal, G. W. & Adler, J. The Journal of General Physiology 62, 203–223, (1973).

28 Lewus, P. & Ford, R. M. Biotechnology and Bioengineering 75, 292–304, (2001).

29 Ahmed, T., Shimizu, T. S. & Stocker, R. Nano Letters 10, 3379–3385, (2010).

30 Boyd, S., Parikh, N., Chu, E., Peleato, B. & Eckstein, J. Foundations and Trends in Machine Learning 3, 1–122, (2011).

31 Ely, B. Methods in Enzymology 204, 372–384, (1991).

